# Proteomic analysis of flagella from *Chlamydomonas* mutants lacking the central pair apparatus reveals loss of radial spoke head proteins

**DOI:** 10.1101/2022.10.02.510551

**Authors:** Kimberly A. Wemmer, Juliette Azimzadeh, Wallace F. Marshall

## Abstract

The flagellar axoneme is a complex protein-based machine capable of generating motile forces by coordinating the action of thousands of dynein motors. A key element of the axoneme is the central pair apparatus, consisting of a pair of microtubules surrounded by additional structures. In an effort to better understand the organization of the central pair, we used 2D DIGE to identify proteins that are depleted from flagella isolated from two different *Chlamydomonas reinhardtii* mutants, *pf15* and *pf18*, that lack the central pair. The set of proteins contained almost no components of the central apparatus. We find that three proteins of the radial spoke head RSP1, RSP9, and RSP10, as well as a number of other protein components associated with the outer doublets, are depleted from flagella of mutants lacking the central apparatus. Two of the other proteins depleted from *pf15* and *pf18* flagella, the microtubule inner proteins (MIPs) FAP21 and FAP161, are missing from the genome of *Thalassiosira*, an organism that lacks a central pair and radial spokes, and RNAi of FAP21 in planaria shows that it has a role in ciliary motility. Based on the depletion of radial spoke head proteins, as well as MIPs and other axonemal components, from flagella lacking the central pair apparatus, we hypothesize that the central apparatus may play a role in scaffolding the assembly or retention of radial spokes and other axonemal structures.

## Introduction

The proper assembly of complex structures within cells is a challenging self-assembly problem. Flagella and cilia are among the most complex structures found in the cell. The internal structure of most motile cilia and flagella consists of nine parallel doublet microtubules arranged around a central pair of singlet microtubules (**Figure 1A**). Force for propulsion is provided by thousands of dynein motors, anchored in two rows along one side of each doublet and organized into multi-headed complexes called the inner and outer dynein arms (DiBella 2001; Yamamoto 2021), that can walk along the microtubule of the adjacent doublet.

**Figure 1.**
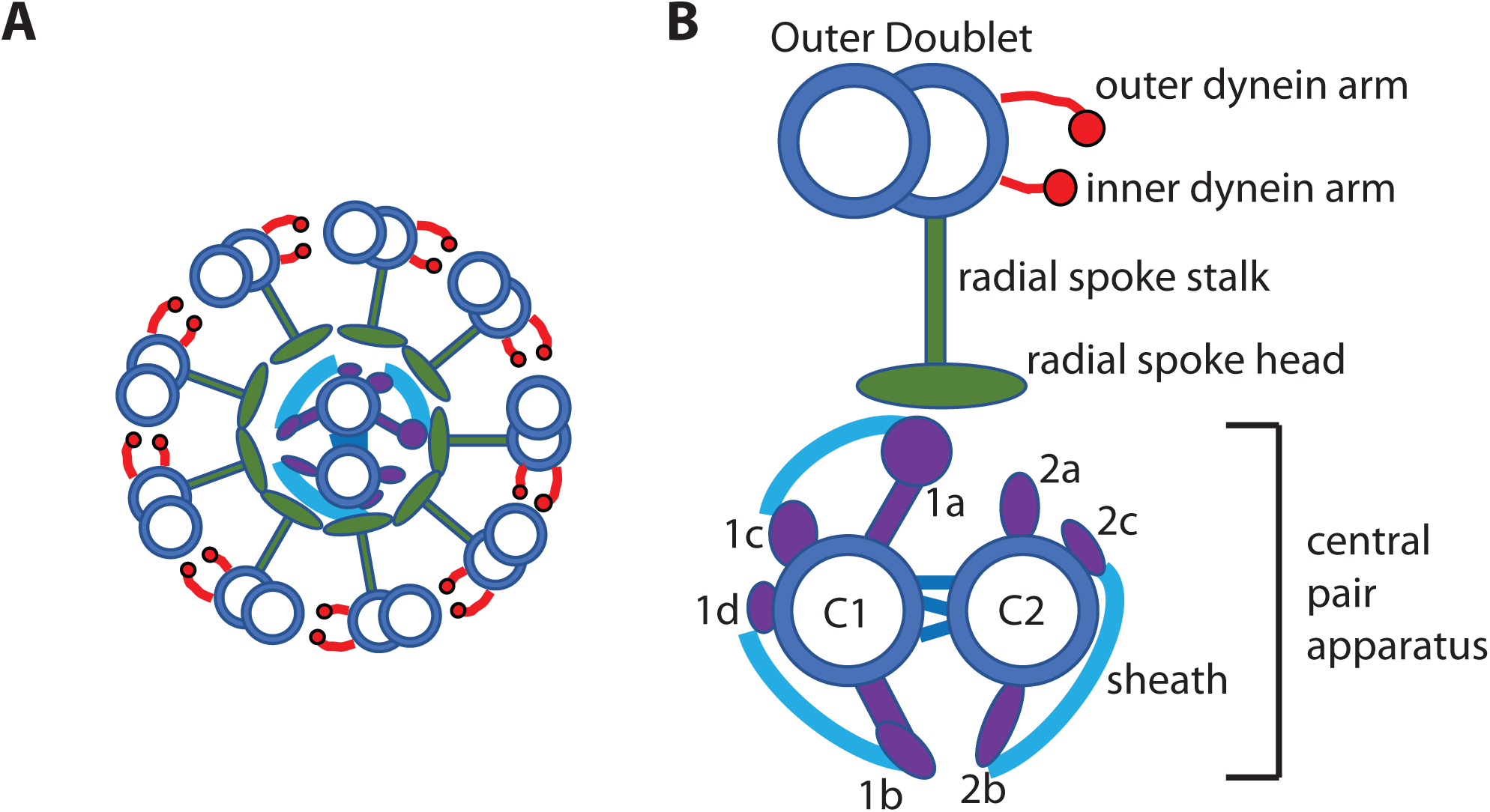
The structure of a flagellum. Microtubules (blue circles) make up the backbone of the structure. (**A**) Cross section of a flagellum, showing nine outer doublets arranged around the central pair microtubules, with radial spokes projecting inward. (**B**) Magnified view of the central pair, consisting of the C1 and C2 tubules with their associated projections (1a-2c, purple) and sheath (light blue); together with a single outer doublet with its radial spoke (green) and inner (ida) and outer (oda) dynein arms (red).

The central pair apparatus is a complex sub-structure within the flagellum, which contains not only a pair of microtubules, C1 and C2, but also an elaborate set of projections (**Figure 1B**). Various projections from the central pair apparatus are responsible for different behaviors, for example, studies in sea urchin sperm and *Chlamydomonas* show that only those dynein arms located near one side of the central pair C1 microtubule are active, while the rest are inactive (Nakano et al., 2003; Wargo and Smith, 2003).

Radial spokes are protein complexes consisting of a stalk anchored on the outer doublet microtubules and extending a head inwards towards the central pair apparatus (Curry, 1993; Diener 2011; Poghosyan 2020). The projections on the central pair interact with the radial spoke head (Kohno et al., 2011), inducing a change in state on the radial spoke. The resulting signaling cascade is not well understood, but likely involves use of secondary messengers, especially calcium, signaling to the dynein regulatory complex located on the A tubule of the outer doublets, which in turn triggers activation of a subset of dyneins (Heuser et al., 2009; Bower 2013; Oda 2015; Gui 2019). In order for all these interactions to occur in their prescribed manner, cells must properly assemble the proteins of the central pair apparatus in the proper structure, correctly aligned with the radial spokes.

Two mutants have been described in *Chlamydomonas, pf15* and *pf18*, which have paralyzed flagella and which lack the organized central pair apparatus structure (Warr 1966; Adams 1981; Witman 1978). The *pf15* strain carries a mutation in the p80 subunit of Katanin, a heterodimeric protein with microtubule-severing activity (Dymek 2004) while the identity of the PF18 gene is not known. Previous studies reported that the flagella of *pf18* cells fixed in situ retain a large amount of electron dense material within their center, where the central pair structure would typically be located (Adams et al., 1981). A similar electron dense core was reported in purified axonemes of *pf15* (Witman 1978). However, when axonemes are isolated from *pf18*, the dense material is lost from the center of the axoneme, suggesting that the proteins present in *pf18* flagella may have been removed during axoneme preparation (Zhao 2019). This fact was used to analyze the proteome of the central pair apparatus by comparing the proteins found in axonemes isolated from wild type versus *pf18* mutant cells (Zhao 2019).

Using a small panel of antibodies directed against selected central pair apparatus proteins, Lechtreck (2013) found that a subset of central pair proteins, normally associated with the C1 microtubule, are specifically retained in *pf15* and *p18* flagella even though they are missing from isolated *pf18* axonemes, while C2 associated proteins were missing from the flagella of the mutants as well as from the axonemes. This was an extremely interesting result, because it implies that central pair apparatus C1 proteins can be imported into flagella, and placed in roughly the right location, but cannot be assembled into the proper arrangement in these mutants. Based on this study, we took an alternative, more systematic approach, in which we identified proteins depleted from isolated whole flagella from *pf15* and *p18* mutants, in the hopes of being able to divide up the central pair apparatus proteins into two groups, one that is retained in flagella and therefore likely to be associated with the C1 part of the complex, and another set depleted from the flagella and therefore likely to be associated specifically with the C2 microtubule. Contrary to our expectation, with a single exception, the proteins that we found to be depleted from *pf15* and *pf18* flagella did not correspond to any of the known central pair apparatus proteins. Instead, we found that a specific set of radial spoke head proteins, RSP1, RSP9, and RSP10, were depleted in both *pf15* and *pf18* flagella, raising the possibility that a correctly organized central pair apparatus is required to scaffold the proper assembly of other axonemal structures such as the radial spokes. Our study also reveals the microtubule inner protein (MIP) FAP21 to be a protein involved in motility and whose presence correlates with the presence of the central pair apparatus.

## Results

### Proteomic analysis of central apparatus mutants

We isolated *Chlamydomonas* flagella, as detailed in Materials and Methods, by subjecting cells to pH shock, and isolating flagella using sucrose cushions to separate flagella from cell bodies. The efficiency of the pH shock and enrichment of flagella in the final preparation were confirmed visually using DIC microscopy, which also allowed for the verification of low amounts of cell body contamination of the preparation. As shown in **Figure 2**, flagella from WT, *pf15*, and *pf18 Chlamydomonas* strains were isolated and differentially labeled, then subjected to 2D DIGE (Viswanathan 2006), in which the proteins of the isolated flagella were separated using 2-dimensional gel electrophoresis after fluorescent labeling. This method provides a quantitative readout of protein differences between two samples by measuring the ratio of fluorescence intensity of each 2D gel spot in both wavelengths. The fold enrichment of individual spots on the gel was determined for each mutant relative to WT samples run on the same gel, and spots with a 1.5 fold change or more in either mutant relative to WT, were excised from the gel and their identity determined using mass spectrometry.

**Figure 2.**
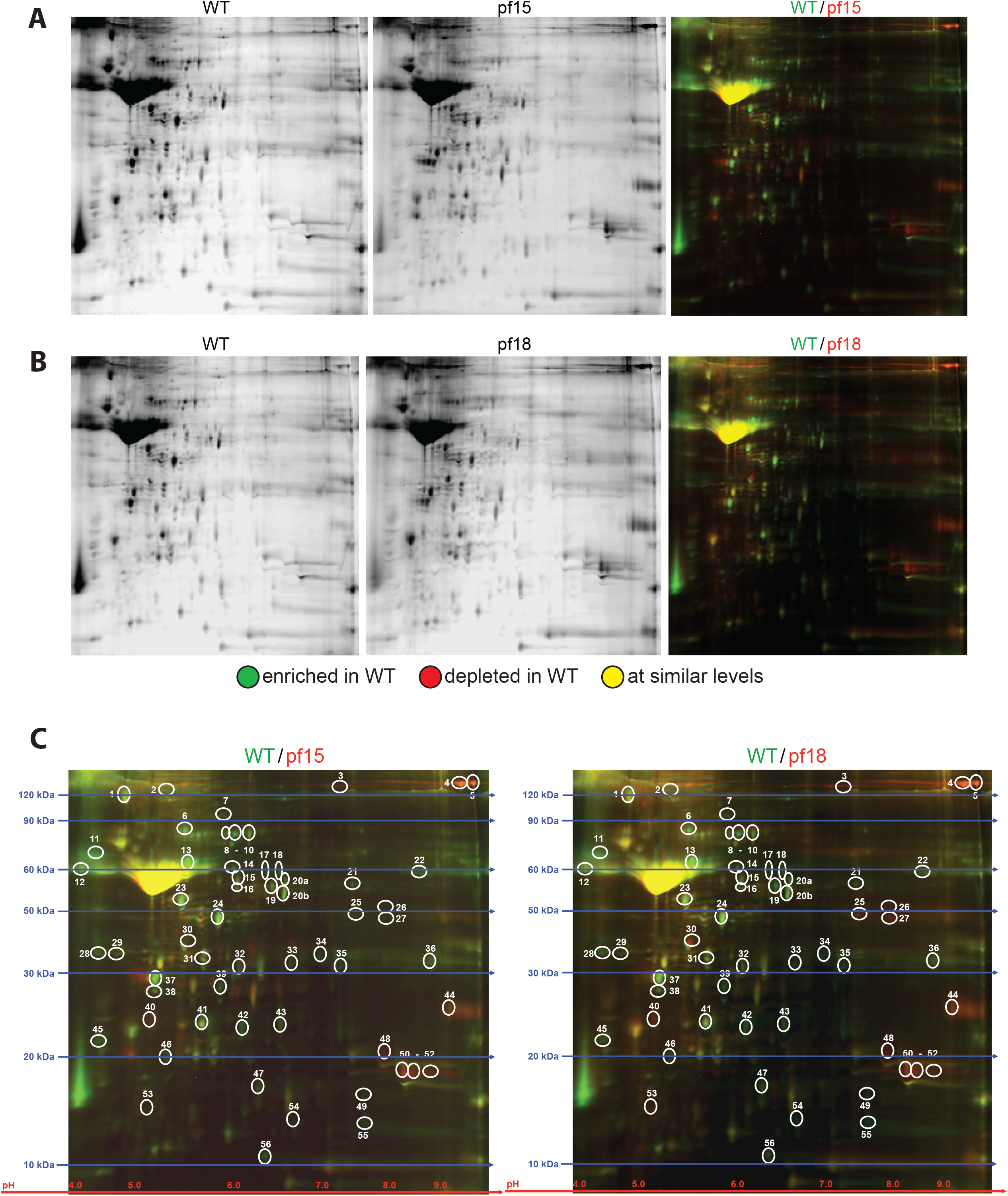
Gel images from 2D DIGE experiment. (**A**) Images of the individual channels of WT and *pf15*, as well as an image of the overlay of the two channels together. (**B**) Equivalent images to (A), but for WT and *pf18*. (**C**) Images showing the comparisons for WT:*pf15* and WT:*pf18*, with circles indicating the spots picked for analysis. Estimated molecular weights (kDa) are indicated by the blue lines on the gel, and the pH range for isoelectric focusing (pH) by a red line at the bottom of the gel.

This analysis provided a list of 57 proteins with differential abundance in the flagella of both mutants (**Table 1**), out of which 42 were depleted in both *pf15* and *pf18* relative to wild type, while 13 were enriched in both *pf15* and *pf18*. Two proteins (FAP155 and RAN1) behaved differently in the two mutants.

**Table 1.**
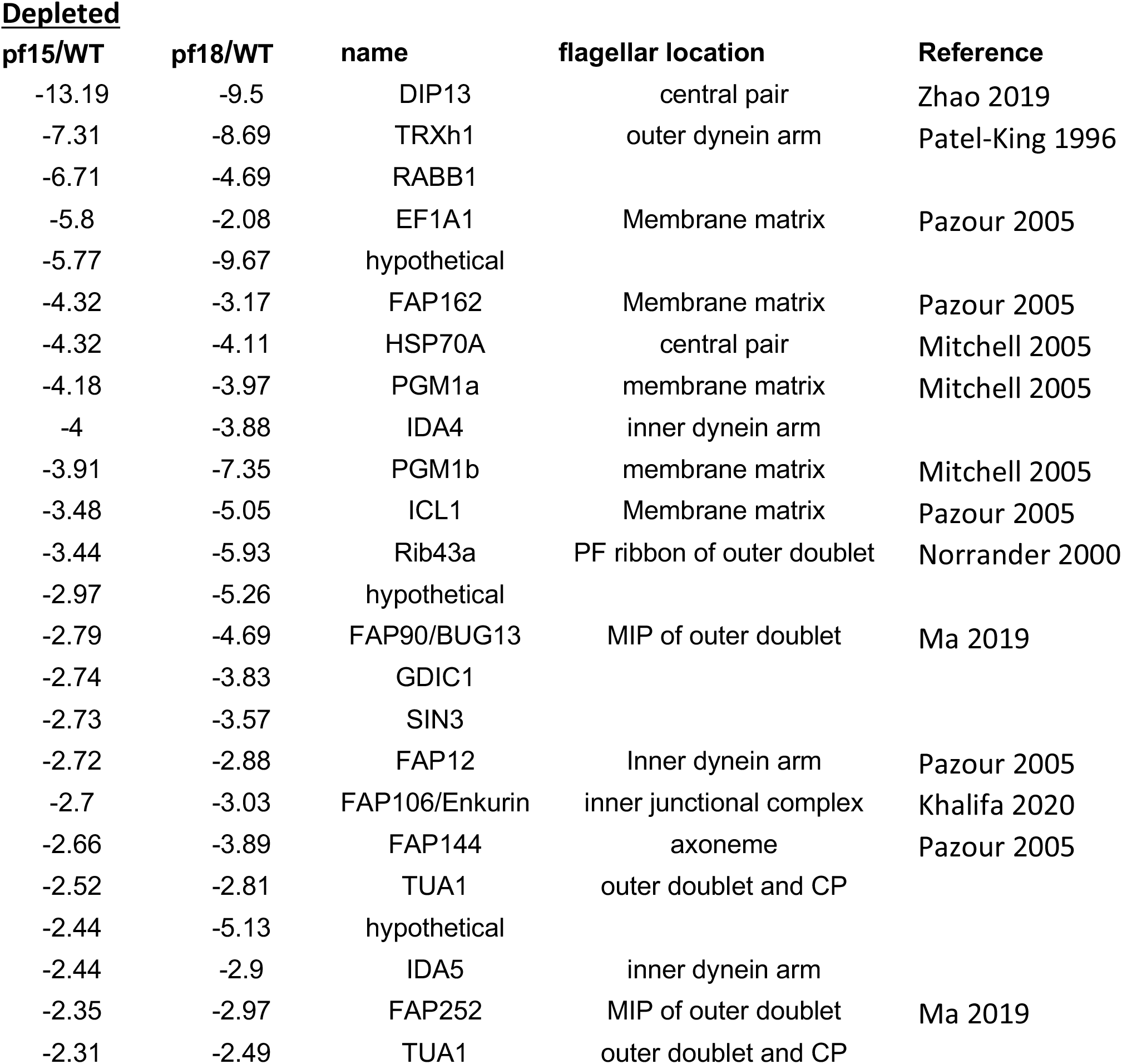
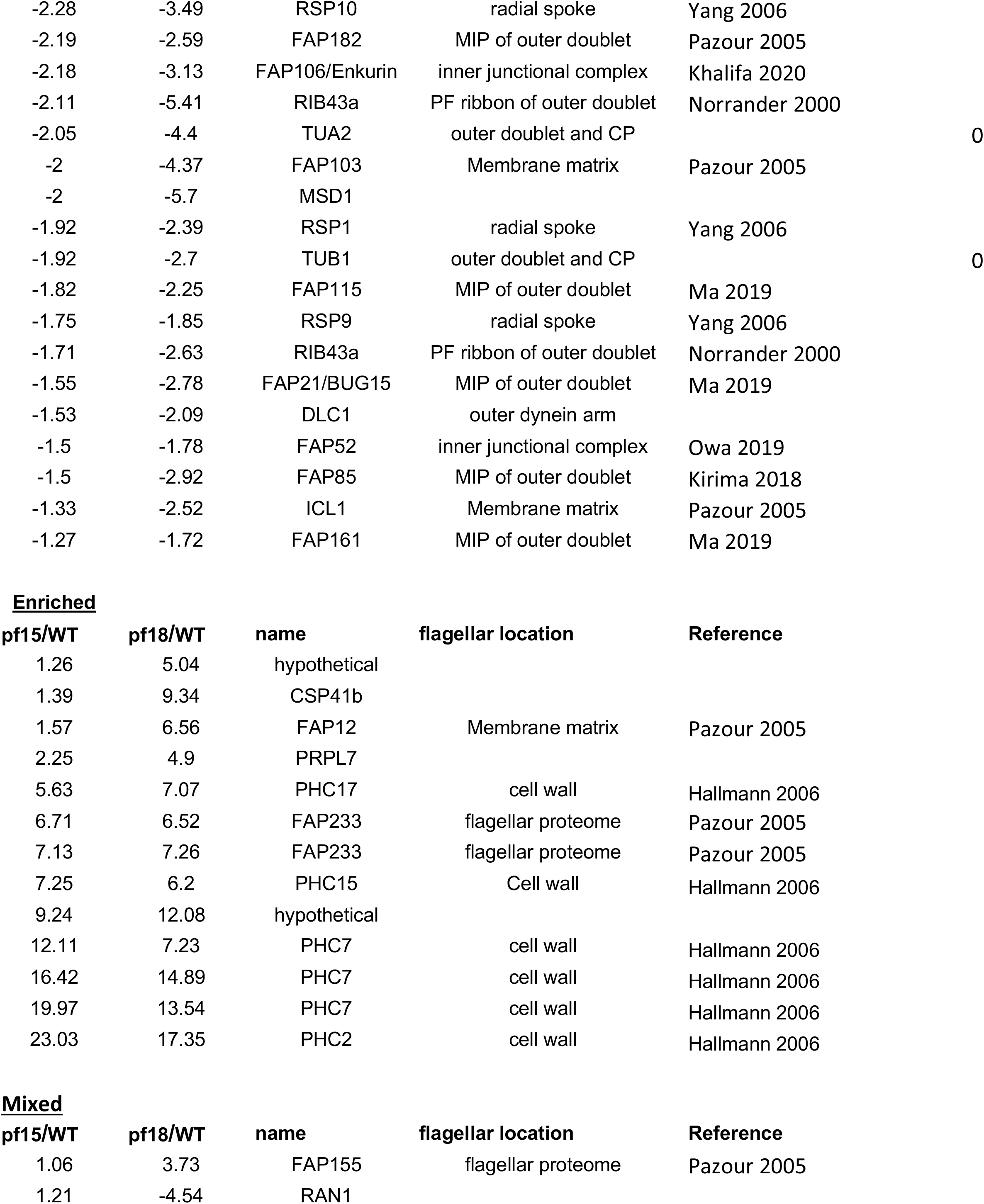
Proteins depleted from both pf15 and pf18, or enriched in both, or showing a mixed effect being depleted from one and enriched in the other. The first two columns give the ratio of fluorescence between each mutant and wild-type flagella, with negative values giving the fold depletion in the mutants and positive values the fold enrichment. Each line represents one spot identified on the 2D DIGE gels. In some cases, the same protein was present in more than one spot, likely due to phosphorylation or other modifications, and in those cases the protein will be listed more than once in the table.

Out of the 42 proteins detected as being differentially depleted in both *pf15* and *pf18* mutant flagella, the majority (32) were known flagellar proteins. Eight proteins were clearly non-flagellar contamination, given that they are either cell wall or chloroplast components, but all of these were among the differentially enriched, rather than depleted, proteins. The degree of enrichment or depletion was highly correlated between the two mutants (r=0.91), suggesting that even though the *pf15* and *pf18* mutations are in different genes, their effects on flagellar protein composition were similar.

Contrary to our initial expectation, the most well-known central pair proteins, for example CPC1, Hydin, PF20/SPAG16, and PF16/SPAG6, were not included in our list. In fact, the only two central pair apparatus proteins identified were Hsp70 (Mitchell 2005), which is involved in many cellular processes, and DIP13, a protein whose expression is upregulated during flagellar regeneration (Stolc 2005), which has been identified as part of the central pair apparatus by a proteomic analysis (Zhao 2019), but which also localizes extensively to the outer doublet microtubules (Pfannemschmid 1993). Thus, neither of the central pair proteins depleted in the mutant flagella proteomes localize exclusively, or even primarily, in this structure. We do however note that DIP13 was the most highly depleted protein in both mutants (**Table 1**).

### Proteins depleted in central apparatus mutants

Among the proteins depleted in *pf15* and *pf18* mutant flagella, three protein components of the radial spoke, RSP1, RSP9, and RSP10. Two of these proteins, RSP1 and RSP10, contain MORN domains. All three of these proteins are part of the radial spoke head (Yang 2006), and it has been speculated that these proteins in particular may take part in direct interactions with the central pair apparatus (Yang 2006). Consistent with this idea, structural studies show that RSP1 and RSP10 form protrusions from the spoke head (Grossman-Haham 2021).

In addition to the radial spoke head proteins, we also found that the *pf15* and *pf18* flagella were depleted for Rib43a, a component of the protofilament ribbon (Norrander 2000; Ichikawa 2019); Fap52 and Fap106, two components of the inner junctional complex of the outer doublet (Owa 2019; Khalifa 2020); FAP85 and FAP115, seven Microtubule Inner Proteins (MIPs) of the outer doublet (Kirima 2018; Ma 2019; Li 2021); and thioredoxin, which is part of the outer dynein arm (Patel-King 1996). In fact, virtually every part of the axoneme except the central pair apparatus is well represented among the proteins depleted from the two ostensible central pair apparatus mutants.

Also depleted was phosphoglyceromutase (PGM), a glycolytic enzyme that is normally found in flagella. Although PGM is in the soluble membrane+matrix fraction (Mitchell 2005), the downstream glycolytic enzyme enolase is part of the central pair apparatus (Michell 2005), possibly suggesting that the central apparatus plays a role in retaining or organizing these enzymes.

### Comparative genomics

As an alternative strategy to determine proteins that might depend on the central apparatus for their presence in flagella, we tested which of the proteins from Table S1 were represented in the genomes of other species with motile cilia, but absent from the genome of *Thalassiosira pseudonana. Thalassiosira* are well-studied marine diatoms, and this particular species was the first eukaryotic marine phytoplankton to be sequenced (Armbrust 2004). Importantly, *Thalassiosira* has flagella that lack central pairs, along with other flagellar structures that are responsible for regulating waveform, including the radial spokes and inner arm dyneins (Heath and Darley, 1972; Manton et al., 1970). This absence of the central pair makes it ideal for this type of comparison - genes for proteins that are lacking in *Thalassiosira* flagella are more likely to either be central pair components, or to rely on the central pair for their assembly into flagella. Among the proteins we have identified as depleted from central apparatus mutant flagella in *Chlamydomonas*, we failed to find orthologs in *Thalassiosira* for only 4: IDA4, FAP21, RSP9, and FAP161. IDA4 and RSP9 are already known as inner arm dynein and radial spoke proteins, respectively. The other two proteins, FAP161 and FAP21, have been identified as a microtubule inner proteins (MIPs) of the outer doublet microtubules (Ma 2019), but no functional information is available about either protein. They are missing from the genomes of species that lack motile cilia but also from *Thalassiosira* (**Figure 3**).

**Figure 3.**
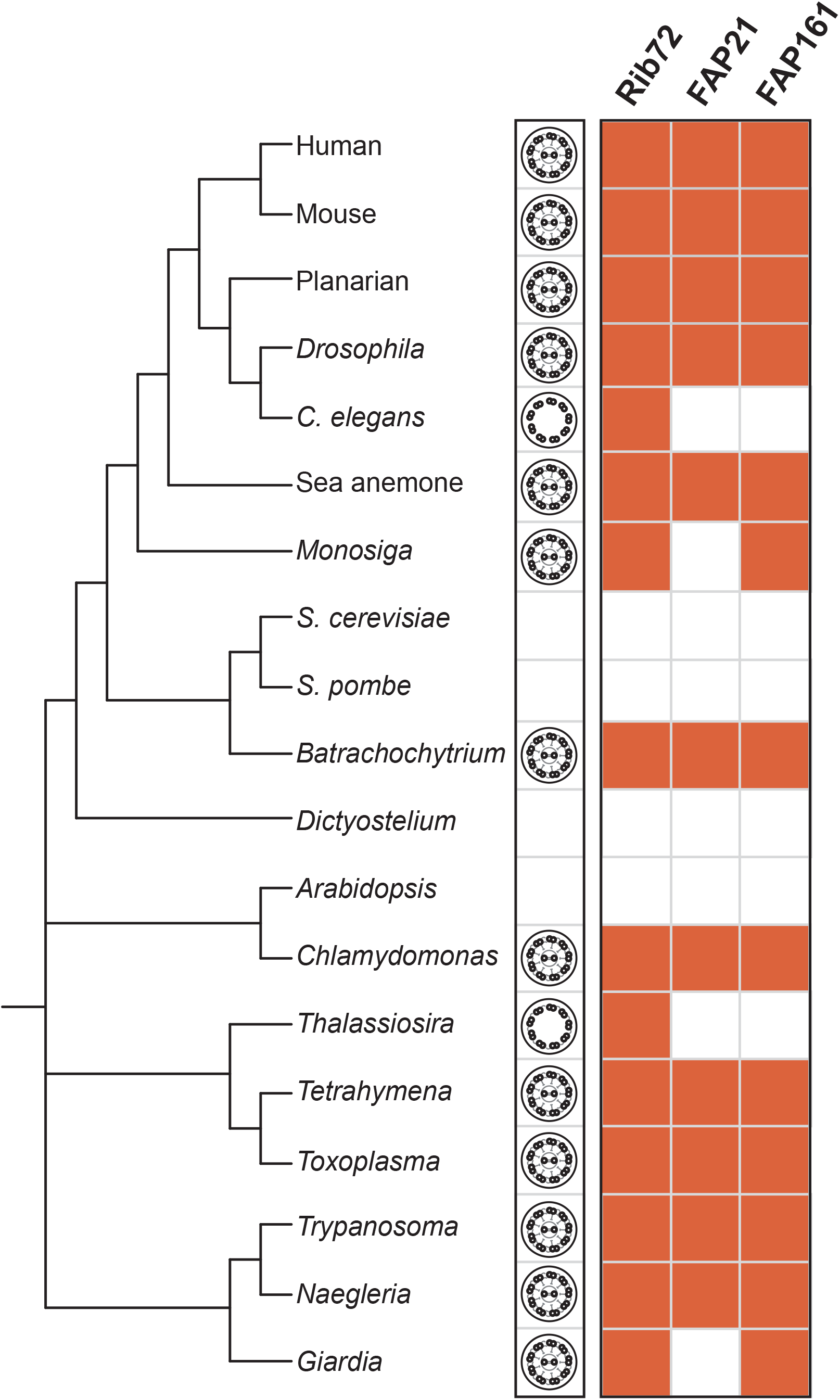
FAP 21 and FAP161 are present only in organisms which have motile cilia with central pairs. This figure is adapted from the Tree of Life project, and is based on the relationships between the different species. The presence or absence of the FAP homologs in each species is indicated with the presence or absence of an orange box. The diagrams on the left column illustrate the axoneme configuration in each species, with an empty box denoting species that lack cilia and flagella entirely. FAP161 and FAP21 appear only in organisms with motile cilia containing central pairs, and are absent in organisms lacking this structure. Rib72 is included as a control; it is a structural component of the A-tubule of the outer doublets. It therefore can be found in all organisms with cilia, but is absent from organisms that do not have cilia or flagella.

### RNAi of FAP21 in Planaria

The absence of FAP21 in *Thalassiosira*, which has motile flagella but lacks the standard machinery for coordinating flagellar motion via radial spokes and a central pair apparatus, suggested the possibility that this protein might be a component of the flagellum involved in regulating waveform or motility. The planarian flatworm *Schmidtea mediterranea* is a rapid and convenient platform in which to test candidate genes for a role in ciliary motility. The organism is covered in cilia, with the cilia on the ventral surface of the worm used to drive motion over a surface, and RNAi is typically fairly robust and easily accomplished in Planaria via injection of double-stranded RNA (dsRNA) (Rompolas et al., 2010). We identified an ortholog of FAP21 in Schmidtea (gene mk4.003874.01.01) and generated constructs to produce dsRNA against this gene.

In Planaria injected with dsRNA corresponding to the homolog of FAP21, we were able to observe a significant motility defect (**Figure 4**). Microscopic analysis indicated that many of the cilia on the ventral surface of the worms subjected to RNAi were still motile, indicating that the RNAi did not cause completely ciliary paralysis. This was expected due to the phenotype we saw - complete paralysis of the cilia is known to result in an inching-worming behavior, which we did not observe. Under the imaging conditions employed, it was not possible to determine whether some cilia might have been fully paralyzed.

**Figure 4.**
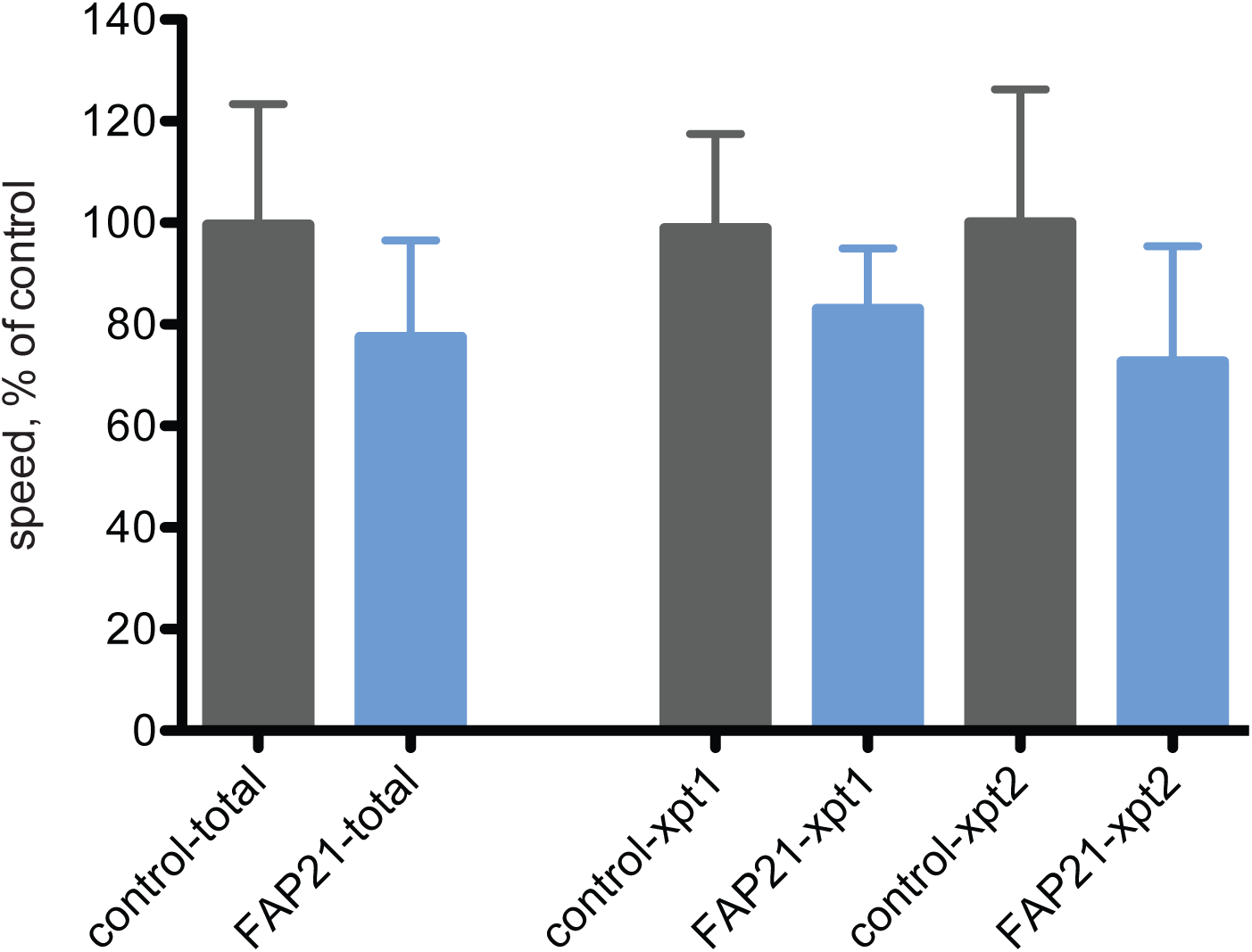
Knock-down of FAP21 impairs cilia-driven motility in Planaria. RNAi knock-down of FAP21 in *S. mediterranea* results in impaired movement of the flatworm. Their motility is reduced by 20% when compared to control worms. This data represents the quantification of two separate experiments, denoted xpt1 and xpt2, with the analysis of all collected data represented in the ‘total’ columns.

## Discussion

### Why weren’t known central pair proteins detected?

Given that our study analyzed proteins depleted from flagella lacking the central apparatus, the obvious expectation would have been that most of the proteins identified would be part of the central apparatus itself, but this was not observed.

One potential explanation for these results could be that the electron dense material noted in the flagella of these mutants is composed of central pair proteins, that were transported to the correct sub-cellular location, but were then unable to assembly into a stable structure that could survive the process of axoneme isolation (Lechtreck 2013). The presence of this material in isolated flagella, which were used in our study, would result in little alteration in their levels when compared to wild type, such that our analysis of differential enrichment would not have detected them. Our results are thus consistent with a picture in which the mutants *pf15* and *pf18* are required for assembling central apparatus proteins into the correct structure, but are not required for the synthesis of these proteins or for their transport or retention in the flagellum. Our results differ from those reported by Lechtreck (2013) who found that C1 associated proteins were retained in *pf18* and *pf19* mutants. One possible difference is that our analysis looked for proteins retained in both *pf15* and *pf18* whereas Lechtreck focused on *pf18* and *pf19* individually. Differences could also simply reflect differences in the detection threshold for 2D DIGE versus Western blotting. Further analysis will be required to establish with certainty the degree to which C1 versus C2 proteins are retained proteome-wide. In any case, our results are broadly consistent with the prior work of Lechtreck in the sense that flagella from mutants lacking an organized central pair apparatus may still contain a large number of the proteins that would normally compose this structure.

However, a more trivial explanation may simply be that because an axoneme only contains a single central pair, but nine outer doublets along with their associated structures (dynein arms, radial spokes, DRC, etc), it is to be expected that central pair proteins will be much fainter on the 2D DIGE gel. Even if the central pair apparatus proteins were technically within a detection threshold, they would tend to be missed in favor of the much more abundant proteins associated with the outer doublets.

The one central apparatus protein that was highly depleted was DIP13, which is a microtubule remodeling protein (Basnet 2018). Although electron microscopy has shown that there is substantial material remaining in the core of the axoneme in the two mutants, the one structure that is certainly missing is the organized pair of microtubules. We hypothesize that DIP13 might be uniquely dependent on these microtubules for its retention inside the axoneme core.

### A scaffolding function for the central apparatus

The present study finds that when the central apparatus is absent, radial spoke head proteins, as well as proteins localizing to the protofilament ribbon and inner junctional complex are depleted, suggesting that the central apparatus is playing a crucial role in supporting the assembly of other structures. We believe that these central apparatus interactions with other axonemal structures may represent a clear example of the general concept of one cellular structure scaffolding another.

We also note that the dependence can go the other way - it has been found that in mutations affecting radial spokes, the central pair apparatus can be affected (Castleman 2009; Jeanson 2015). Such complex interactions suggest that phenotypes resulting from mutations in one part of the axoneme can have effects that ramify to other parts, possibly contributing to the great pleiotropy seen in human ciliary disease mutations.

With respect to the radial spoke head proteins depleted in pf15 and pf18, it is interesting to note that RSP1 and RSP10 had previously been proposed to interact directly with the central pair apparatus, based on the presence of interaction motifs found in these proteins (Yang 2006).

## Materials and Methods

### Chlamydomonas strains used in this study and culture conditions

Strains cc125 (WT), cc1033 (*pf15)*, and cc1036 (*pf18)* were used in the 2D DIGE analysis. All strains were obtained from the *Chlamydomonas* Center (www.chlamy.org). Strains were maintained on TAP plates, then were grown in liquid TAP media in constant light at 21C with agitation (5mL cultures in 18mm test tubes were grown on roller drums, 100-200 mL cultures were grown on an orbital shaker running at 150 rpm, and large 1L cultures were grown with gentle stirring and bubbling with house air).

### Isolation of Flagella from Chlamydomonas

1-3 small cultures of 120 mL were started with half a loop’s worth of cells from TAP plates, then were grown for 3 days as described above. 100 mL of each of these cultures were used to start a 1 L TAP culture in a 2 L flask which was grown for 5 to 7 days. Cultures were checked visually for the presence of flagella. Each 1 L culture was spun down in 1 L bottles at 1500xg in a Beckman Coulter Avanti J-20XP centrifuge in a JLA 8.1000 rotor for 10 minutes (min) at 19-21C with a low brake. The cells were resuspended in 150 to 200 mL TAP media, transferred to a Corning 250 mL centrifuge tube (430776) and then spun again at ∼1000xg (2000rpm in a Beckman Coulter Allegra 6 with a GH3.8 rotor) for 10min at RT with no brake. Cells were re-suspended in 80mL TAP and transferred to a flask, where they were allowed to recover for 2 hours at RT in the light with gentle stirring and bubbling. These cultures were then pH shocked in the following way: a calibrated probe from a pH meter was inserted into the culture, at 0min 0.5 M acetic acid was added until the pH of the culture was between 4.5 to 4.6 typically within 30 seconds of first adding the solution. At 1.5min, 0.5 M KOH was added until the culture reached a pH of between 7 and 7.2, again typically within 30 seconds, then 100 μL of Sigma protease inhibitor cocktail (P8340) and 100 μL of 100 mM of EGTA, pH7.5 were added to the culture, which was then transferred to a 250mL tube and placed on ice. The efficiency of the pH shock was verified visually. The rest of the protocol was completed on ice. The culture was spun at ∼1000 xg (2000 rpm GH3.8 rotor in an Allegra 6R centrifuge) for 10 min at 4C with no brake. The supernatant was removed and transferred to a new tube, and the cell bodies were discarded. 50 mL of 25% sucrose in TAP with red food coloring, to allowed it to be seen, was pipetted beneath the supernatant as an underlay, and the tube was spun at ∼2500xg (3300 rpm GH3.8 rotor) for 10 min at 4C with no brake. The entire supernatant and the interface between the two layers was collected, transferred to a new tube, and the entire underlay process repeated. The entire supernatant and interface was again collected, transferred to a new tube, and placed on ice. 35 mL of this supernatant at a time was transferred to a 50 mL Nalgene polycarbonate centrifuge tube (3117-0500) and spun at ∼16000xg (10000rpm in a HB-4 rotor in a Sorvall RC5C centrifuge) for 20 min at 4C. The supernatant was discarded, and the next aliquot placed in the tube and spun. This was repeated until the entire sample was pelleted. The pellet was re-suspended in 1mL Applied Biomic’s Cell Washing Buffer (10 mM Tris-HCl, 5 mM magnesium acetate, pH 8.0) and transferred to a 1.5 mL microfuge tube. The quality of the isolation was assayed visually, checking for high concentrations of flagella with low amounts of cell body debris. The isolated flagella were pelleted one final time at ∼20000xg (14000rpm in an Eppendorf 5417C centrifuge) for 20 min at 4 C, the supernatant aspirated, and the pellet frozen in liquid nitrogen, then transferred to a -80 C freezer.

### 2D DIGE and protein identification

Sample labeling, 2D DIGE, image analysis, spot analysis using DeCyder software, excision from gel, and identification using mass spectrometry were all performed by Applied Biomics, Hayward, CA (http://www.appliedbiomics.com).

### cDNA cloning

FAP21 cDNA was obtained by RT-PCR on total RNA isolated using TRIzol (Invitrogen, 15596-026) according to the manufacturer’s instructions. For Planaria, the RNA was isolated from the asexual strain, the cDNA sequences for our genes of interest amplified using the primers listed below, and the sequences were cloned into pPR-T4P, a modified pDONR-dT7 in which the gateway cloning site was replaced by a ligation-independent cloning site (kind gift from J. Rink). The primers for FAP21 were as follows:

Forward 5’ CATTACCATCCCGGCCTAATCATGTGTTTGGAAC 3’

Reverse 5’ CCAATTCTACCCGGCCAGTTAAAGCTGTCAGTAAA 3’

### RNAi via dsRNA injection

Sense and antisense DNA templates for *in vitro* transcription were obtained by amplifying sequences cloned into pPR-T4P using the following primer pairs:

*Sense* F: AACCCCTCAAGACCCGTTTAGA and R: GAATTGGGTACCGGGCCC;

*Antisense* F: CCACCGGTTCCATGGCTAGC and R: GAGGCCCCAAGGGGTTATGTG

dsRNA was synthesized using the T7 Ribomax Express RNAi system (Promega, Madison, WI). For RNAi experiments, 10 ∼ 0.8-1cm long planarians were injected 3 consecutive days with 20-50 ng dsRNA per injection using a Nanoject II injector (Drummond Scientific, Broomall, PA). The animals were then amputated from their head and tail at day 4. An additional injection was performed at day 11 and the flatworms were amputated again pre- and post-pharyngically at day 12. Phenotypes were assessed on regenerating heads, tails and trunks at day 21 (10 days after the second amputation).

### Imaging of Planaria

Live images were acquired using a Stemi 2000C stereomicroscope equipped with an AxioCam MRc digital microscope camera (Carl Zeiss MicroImaging, Thornwood, NY). Planarian locomotion speed was determined using ImageJ software, by measuring the distance covered by a single worm between frames separated by known time intervals.

## Acknowledgments

We thank members of the Marshall lab for advice and interesting discussions, and members of the Sanchez Alvarado lab for advice about Planarian experiments. This work was supported first by NIH grant R01 GM097017 and subsequently by R35 GM130327.

## Note

This is a preprint and is not peer reviewed

